# Vertebrate Biodiversity via eDNA at the air-water interface

**DOI:** 10.1101/2025.06.13.659655

**Authors:** Yin Cheong Aden Ip, Pedro FP Brandão-Dias, Gledis Guri, Elizabeth Andruszkiewicz Allan, Ryan P. Kelly

## Abstract

Although aquatic, aerial, and terrestrial habitats are often treated as separate ecological systems, these environments exist along a continuum of connectivity: flows of biomass and energy routinely create linkages across water and air, blurring traditional boundaries. Here, we use eDNA sequencing to illustrate and quantify the movement of trace genetic information between water and air. We collected 27 paired air-and-water samples from two urban–wildland interface sites using passive air sampling and active water filtering. Metabarcoding with the MiFish-U 12S marker recovered 35 vertebrate taxa, 40% of which were detected in both water and air, ranging from the strictly aquatic salmon to wholly terrestrial cottontail rabbit. This reciprocal relationship suggests that eDNA pools form within each partition (water or air), with the probability of transfer governed by DNA concentration. In this view, detecting aquatic eDNA in the air is not contamination or stochastic noise but an expected, repeatable phenomenon. Logistic models confirm that higher abundance in one medium predicts spillover into the other. For instance, peaks in Coho and Chinook salmon eDNA align within 24 hours, demonstrating that passive air sampling reflects the temporal abundance trends of the most common aquatic species. In contrast, low-read-abundance taxa appear only sporadically, implying that rare detections are the first to drop out of cross-medium transfer and therefore demand intensified sampling in their primary habitat. Together, our findings bridge the conventional separation of aquatic and airborne eDNA and establish a unified, non-invasive framework for holistic vertebrate monitoring at the land–water interface. This approach offers transformative potential for conservation, invasive-species early warning, and One Health surveillance.

## 1. Introduction

Multicellular species continually shed genetic traces into their surroundings, leaving environmental DNA (eDNA) in water and air. For the most part, eDNA surveys treat these two reservoirs as separate, as researchers assume the water samples predominantly yield aquatic species, and air yields primarily terrestrial and airborne taxa. For instance, water samples routinely reveal fish, amphibians, and macroinvertebrates (Deiner et al., 2017; Valentini et al., 2016), and air filters recover bird, mammal and plant DNA from the atmosphere (Johnson et al., 2023; Lynggaard et al., 2022). As a result, “exogenous” signals, which are DNA from organisms not resident in the sampled medium, are often dismissed as incidental or ignored.

In reality, physical and biological processes routinely transport genetic material across environmental boundaries. Rainfall and overland flow can carry terrestrial DNA into streams; surface splashes and bubble-burst aerosolization propel waterborne DNA skyward; windblown dust and settling particles shuttle DNA molecules among air, and water (Allen et al., 2022; Evangeliou et al., 2020). Consequently, trace signatures of nonresident taxa are expected to appear in eDNA surveys. In this context, we define “exogenous eDNA” as genetic material detected in a medium where the source organism is not resident. For instance, terrestrial mammals in water samples and aquatic fish DNA in air filters (Ip et al., 2025a; Darling et al., 2021; Martel et al., 2021). Such exogenous detections can arise not only from abiotic transport but also from biotic and human vectors (Barnes & Turner, 2016). Predators and scavengers may carry prey DNA across habitats, and human activities such as bait fish disposal, livestock grazing near banks, or urban runoff of food waste, can introduce non-resident DNA into aquatic and airborne environmental samples (Klepke et al., 2022). Because eDNA studies typically target species native to the sampled medium, such cross-compartment signals are difficult to interpret without additional ecological context. Incorporating background ecological knowledge and land-use history is therefore essential to distinguish genuine continuum signals from vector-driven deposits and to interpret unexpected detections responsibly.

Our recent quantitative analysis of cross-medium genetic signal transfer showed that spawning salmon release detectable aquatic eDNA into the atmosphere, and that simple passive air samplers recover fish-derived DNA sequences in direct proportion that tracks abundance alongside visual fish counts (Ip et al., 2025a). Meanwhile, multiple prior airborne eDNA studies in purely terrestrial contexts have demonstrated that passive and pump-assisted air collectors reliably recover resident species’ DNA, from mammals, birds, and plants, carried on ambient aerosols (Clare et al., 2022; Lynggaard et al., 2022; Métris & Métris, 2023; Tournayre et al., 2025; Jager et al., 2025). Together, these complementary lines of work suggest two key points: (i) air sampling can recover large amounts of native eDNA from terrestrial species, and (ii) aquatic organisms can also contribute exogenous eDNA to the same airborne “cloud” of genetic signals. Taken together, this evidence suggests cross-medium detections should not be random scattering of fragments but instead reflect an underlying, abundance-dependent process.

We therefore hypothesize that eDNA persists in pools within each environmental partition (water or air), and that the probability of transfer from one partition to another depends on the concentration of species’ DNA in the source partition, whereby molecules at greater concentration in one partition are more likely to move across partitions. Under this mechanistic model, cross-medium detections should follow predictable, abundance-driven patterns rather than being mere artifacts or random noise. To test the hypothesis of an eDNA continuum with bidirectional transfer between water and air, we address four interrelated questions: (1) Continuum authenticity: Do detections of aquatic species in air (and terrestrial species in water) reflect genuine aerosolization and deposition rather than methodological contamination? (2) Abundance dependency: Does a species’ relative abundance or biomass in its native pool predict its likelihood of appearing in the alternate pool? (3) Temporal synchrony: If water and air truly form a continuum of eDNA signals, will dominant taxa exhibit synchronous temporal trends across both reservoirs? (4) Sampling effort: Because rare species shed little DNA, how much sampling effort is required to recover faint signals reliably in both media?

Answering these four questions on continuum authenticity, abundance dependency, temporal synchrony, and sampling effort, will establish the pillars of the emerging bidirectional eDNA Continuum Paradigm. To address them, we conducted the first side-by-side metabarcoding survey of vertebrate eDNA at the air–water interface. Over seven consecutive weeks, we deployed paired open-water trays for passive air sampling and active river-grab filters for water sampling at two urban–wildland interface sites. By comparing taxon co-occurrence, modeling cross-medium detection probabilities, and examining time-series alignment, we evaluate whether water and air truly function as a unified genetic reservoir. This framework lays the groundwork for a unified, noninvasive approach to survey vertebrate life at the land–water interface, with applications in conservation, invasive-species early warning, and One Health surveillance.

## 2. Materials and Methods

### 2.1 Study sites and sampling design

Between 26 August and 18 November 2024, we conducted 17 overnight sampling trips at two urban–wildland interface sites along Issaquah Creek, Washington, USA. Confluence Park (47.535711° N, –122.039819° W) is a public riparian area with mixed deciduous and coniferous vegetation and intermittent pedestrian traffic. Issaquah Hatchery (47.529501° N, –122.039133° W) is ∼2 km upstream of Confluence Park, and comprises managed salmon-rearing ponds bordered by secondary forest. We scheduled sampling to encompass the October 2024 peak in coho salmon (*Oncorhynchus kisutch*) spawning, based on hatchery visual counts and weekly escapement reports. On each trip, we deployed an open-water tray sampler for passive airborne eDNA at ∼ 09:00 h and retrieved it ∼ 24 h later, immediately followed by collection of a matched river-grab water sample.

### 2.2 River water sample collection

At the start of each trip, we collected 3 L water samples from the surface of the river using sterile 3 L Nalgene Cantene carboys. Water sampling location was immediately adjacent to air sampler locations. Each 3L sample was subdivided into three 1L aliquots, and each aliquot was filtered on site through a 5.0 µm Sterlitech MCE filter (Sterlitech, Kent, WA, USA) via the Smith-Root Citizen Science vacuum pump (Smith-Root, Vancouver, WA, USA). Filter membranes were placed in 1.5 mL DNA/RNA Shield (Zymo Research, USA), transported at room temperature, and then frozen at –20 °C within four hours. Of the three 1L replicates, only one was used for metabarcoding. One field blank (1 L Milli-Q water) was processed at the beginning of each trip through a fresh tubing to check for contamination. All sampling gear was decontaminated with 10% bleach and rinsed with deionized water between uses.

### 2.3 Passive airborne sampling

At each site, a polypropylene tray (25 × 30 × 10 cm; surface area ≈ 750 cm²) was filled with 2 L of molecular-grade deionized water and placed ∼ 1 m above the river water and ∼ 5 m inland away from the riverbank at Confluence Park (from a tree branch), and ∼ 3 m above the water at the hatchery’s fish ladder (from railings). Upon retrieval, tray water was immediately filtered in the field through a 5.0 µm Sterlitech MCE filter using a Smith-Root Citizen Science peristaltic pump (see “River-water grab samples” below). The resulting filter membranes were then treated identically to those from water samples (see above).

To facilitate cross-medium comparisons, we note that each water sample comprised 1 L of river water, whereas each air sample collected all eDNA settling onto a 750 cm² water surface over ∼ 24 h. We did not directly measure the volume of air passing over the tray or quantify an exact deposition rate, so it is not yet possible to translate the tray’s collected material into an equivalent air-volume metric. Consequently, any direct comparison of detection probability between air and water must be viewed qualitatively: a fully quantitative “reads per unit effort” framework would require normalizing air-trap deposition to an estimated air volume (or water volume).

### 2.4 Laboratory processing and sequencing

All molecular work was conducted in a 10% bleach and UV-treated PCR-clean hood to minimize contamination. Frozen filters and tray membranes were thawed on ice, and 300 µL of the DNA/RNA Shield preservative was used directly for extraction with the QIAgen DNeasy Blood & Tissue Kit, following manufacturer protocols but omitting any prior proteinase-K digestion. Extracts were quantified via Qubit High-Sensitivity assays and stored at –20 °C until amplification. We targeted a ∼170 bp fragment of vertebrate mitochondrial 12S using only the MiFish-U primer set (Miya et al. 2015), with the following sequences: MiFish-U-F: 5′-GCCGGTAAAACTCGTGCCAGC-3′; and MiFish-U-R: 5′-CATAGTGGGGTATCTAATCCCAGTTTG-3′.

To enable multiplexing and error-robust sample assignment, 96 custom 14 bp molecular identifiers (UMIs) were appended to the 5′ end of each primer (Ip et al., 2025b). UMIs were designed via a Python script to include fixed “CC” on both ends flanking a 10 bp ATG-rich core, with a minimum Hamming and Levenshtein distance of three between any pair. Each 20 µL PCR contained 10 µL Phusion HF Master Mix (ThermoFisher), 0.6 µL DMSO (3% v/v), 0.5 µg µL⁻¹ BSA, 0.5 µL of each UMI-tagged primer (10 µM), 3.9 µL molecular-grade water, and 5 µL extract. Thermocycling was: 95 °C for 5 min; 35 cycles of 95 °C for 30 s, 60 °C for 30 s, 72 °C for 45 s; and a final extension of 72 °C for 5 min. Duplicate reactions per sample were pooled and purified with AMPure XP beads (0.7×).

Libraries were prepared with the Oxford Nanopore SQK-LSK114 ligation kit following standard protocols, then loaded onto R10.4.1 MinION flow cells for 48 h runs on a MK1B device using MinKNOW v22.12.5.

### 2.5 Bioinformatics pipeline

Nanopore bioinformatics processing followed pipelines developed in Ip et al. (2025b). Raw POD5 files were basecalled with Dorado v0.9.1 in Super-Accurate (SUP) mode on an MSI Raider 18 HX laptop outfitted with an NVIDIA RTX 4090 GPU. The resulting FASTQ reads were demultiplexed, primer and UMI-trimmed using ONTBarcoder2.3 (Srivathsan et al. 2024) with a minimum read length of 200 bp and allowing up to two mismatches per UMI tag. This step also splits self-ligated concatemer reads, ensuring high-confidence assignment of each sequence to its originating sample.

Following demultiplexing, we performed within-sample dereplication and clustering using VSEARCH (Rognes et al. 2016). First, each sample’s FASTQ was converted to FASTA and dereplicated at full length, retaining unique sequences of at least one copy. These dereplicated reads were then clustered at 98% identity to collapse sequencing noise and generate representative centroids for each operational taxonomic unit (OTU). We next computed a SHA-1 hash for each centroid sequence and assembled an OTU table of raw read counts per sample, facilitating downstream filtering and normalization.

To refine this initial OTU table, we applied the LULU curation algorithm (Frøslev et al. 2017) in R, which identifies and merges artifactual OTUs based on sequence similarity and co-occurrence patterns. Using default settings (minimum match = 84%, minimum relative co-occurrence = 0.95), LULU removed OTUs likely arising from PCR or sequencing errors, yielding a curated table of high-confidence centroids. We then reconstructed a filtered OTU count matrix by retaining only those centroids that passed LULU curation.

Surviving centroids were subjected to taxonomic assignment via BLASTn against the NCBI eukaryote “nt” database, as of January 2025. We ran BLASTn (version 2.15.0+) with parameters set for high stringency (≥ 96% identity, word size = 30, e-value ≤ 1e-40, maximum 50 targets) and output formatted to capture scientific and common names, alignment metrics, and taxonomic identifiers. This BLAST output provided a pool of putative taxonomic hits for each centroid.

To derive the most likely taxonomic identity and rank for each sequence, we employed TaxonKit (Shen and Ren, 2021) to compute the lowest common ancestor (LCA) across all top identity BLAST hits per hash. The resulting LCA table was merged back with BLAST metadata to produce a final annotated OTU dataset. Lastly, we conducted manual curation of taxonomic assignments. Each LCA-annotated OTU was manually evaluated for ecological plausibility within the Issaquah Creek watershed. We collapsed several ambiguous records to genus level (e.g., *Cottus* sp., *Microtus* sp.) where species-level resolution exceeded the marker’s discriminatory power or lacked local reference sequences. Singleton OTUs (those with only one read across all samples) were removed as probable artefacts. A small number of salmonid sequences that persisted despite these filters, and that only appeared alongside high-abundance salmon in the same samples, were deemed residual sequencing errors and manually excised. The final dataset comprised 39 vertebrate taxa confidently detected across 27 environmental DNA samples.

### 2.6 Statistical analyses

All statistical analyses were carried out in R version 4.4.1. We began by assessing taxonomic overlap between waterborne and airborne eDNA: raw read counts for each species were converted to presence–absence, and the numbers of taxa unique to each medium or shared across both were depicted in a chord diagram (*circlize*; Gu et al., 2014), with ribbon thickness proportional to log₁₀(reads + 1) and wedges colored by habitat affinity (aquatic, amphibious, terrestrial).

Since our MinION workflow did not include any pre-sequencing normalization of library concentrations – as is common in Illumina workflows – higher raw read counts correspond to higher template concentrations across samples (unlike in Illumina datasets). To test whether the distribution of taxa differed more than expected by chance across habitat groups and media, we again converted read counts into presence–absence for air and water. Next, we grouped species by habitat affiliation (Aquatic, Amphibious, Terrestrial) and tabulated, for each habitat, how many species were detected in air versus in water. This yielded a 3 × 2 contingency table (rows: Aquatic, Amphibious, Terrestrial; columns: “Detected in Air” vs. “Detected in Water”). We then applied Pearson’s Chi-square test to evaluate whether habitat affiliation was associated with differences in detection frequency between air and water, with significance assessed at p < 0.05.

Next, for exogenous DNA transfer, we quantified the ease with which DNA “spills over” from one medium to the other by fitting binomial generalized linear models – that is, we are interested in the probability of detection in one medium (air or water) as a function of its abundance in the other medium. For each taxon *i* detected in sample *j* in medium *m* where *m* ∈{*w, a*} (*w* for water and *a* for air), we modeled the probability of its detection *p* as a function of the log_10_ +1 reads (R) in the opposite medium *k*, where *k* ∈{*w, a*} \{*m*} (i.e., if *m=w* then *k=a* for air and vice-versa) using a logistic regression with intercept β0 and slope β1, as

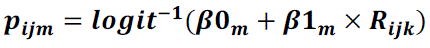

Models were fitted in R using glm(Y ∼ R, family = “binomial”), where y is the 0 or 1 response variable. The resulting logistic curves illustrate how increasing abundance in the source medium predicts cross-medium detections. Since water sampling filtered 1 L per sample while air sampling collected whatever settled onto a 750 cm² tray over 24 h, our logistic-regression curves reflect two very different sampling substrates. Ideally, we would convert air-trap deposition into an equivalent air volume to standardize “reads per unit effort.” In the absence of that normalization, cross-medium detection thresholds should be interpreted qualitatively rather than as absolute concentration comparisons.

To examine temporal concordance between media, we calculated an eDNA-index for each taxon in each sample (Kelly et al., 2019): species read counts were divided by the sample’s total reads to yield proportions, then normalized by each taxon’s maximum proportion (across all samples) to scale the index from 0 to 1. We plotted a time-series of this index for six focal species that represent aquatic, amphibious, and terrestrial guilds, visualizing whether peaks and troughs align across air and water qualitatively. To quantify air–water synchrony for our six focal species, we reshaped each taxon’s eDNA-index time-series into paired columns (index_air and index_water), filling non-detections with zeros, then for each species fit a linear regression of index_air against index_water and recorded the resulting R² as a measure of concordance.

We further explored how detection reliability varies with sampling effort by computing, for each species and medium, the proportion of samples in which it was detected (“detection frequency”) and plotting those frequencies against the square root of total reads per species. This analysis illustrates the diminishing returns of additional sampling for common taxa versus the high effort required to capture rare, stochastic signals.

Finally, to characterize community-level patterns, we generated both Jaccard (presence– absence) and Bray–Curtis (on log10-transformed read counts) dissimilarity matrices and performed non-metric multidimensional scaling in *vegan* (Oksanen et al., 2013) to visualize sample clustering by medium and site. We then used PERMANOVA (adonis2, 999 permutations) to partition the variance in community composition attributable to sampling medium and location.

All visualizations were produced with *ggplot2* and assembled into multi-panel figures using *cowplot*.

## 3. Results

### 3.1 A bidirectional eDNA continuum

We conducted 17 sampling trips over seven weeks, yielding 27 eDNA collections. These comprise 14 airborne (open-water trays) and 13 waterborne (river-grab filters), which together generated 254 252 high-confidence reads after OTU curation (Supplementary Table S1). Across all samples, we detected 35 vertebrate taxa after excluding four putative “food” items Northern Anchovy (*Engraulis mordax*), Pacific Sardine (*Sardinops sagax*), European Sprat (*Sprattus sprattus*), and Wild Boar/Domestic Pig (*Sus scrofa*) (Table 1)—none of which could possibly occur in a freshwater system at Issaquah Creek. Their presence likely reflects human or bait inputs rather than genuine watershed biodiversity. Of these, 14 species (40%) were detected in both water and air (e.g., Coho salmon *Oncorhynchus kisutch*, Mallard *Anas platyrhynchos*, North American Beaver *Castor canadensis*, Domestic Dog *Canis lupus familiaris*), 14 (40%) appeared only in water (e.g., Western Brook Lamprey *Lampetra richardsoni*, Three-spined Stickleback *Gasterosteus aculeatus*), and 7 (20%) were detected only in air (e.g., Pacific Treefrog *Pseudacris regilla*, Song Sparrow *Melospiza melodia*) (Fig. 1, Table 1). The chord diagram illustrates a bidirectional continuum: blue (water) and gold (air) ribbons interweave into a dense web of shared taxa, while the outer arcs highlight each medium’s unique signals and species’ habitat types. Ribbon widths are scaled to log₁₀(reads + 1) and are dominated by a handful of abundant aquatic species (notably the salmon *Oncorhynchus* spp.), which together contribute a substantial fraction of the total eDNA flux in both air and water. Using only presence/absence, a Pearson’s chi-square test on a 3 × 2 table of habitat affinity (Aquatic, Amphibious, Terrestrial) versus detection medium (air vs. water) found no significant association (χ² = 1.57, df = 2, p = 0.46), indicating that, when treating each species as simply “present” or “absent,” the proportion detected in air versus water does not differ by habitat class.

**Figure 1.**
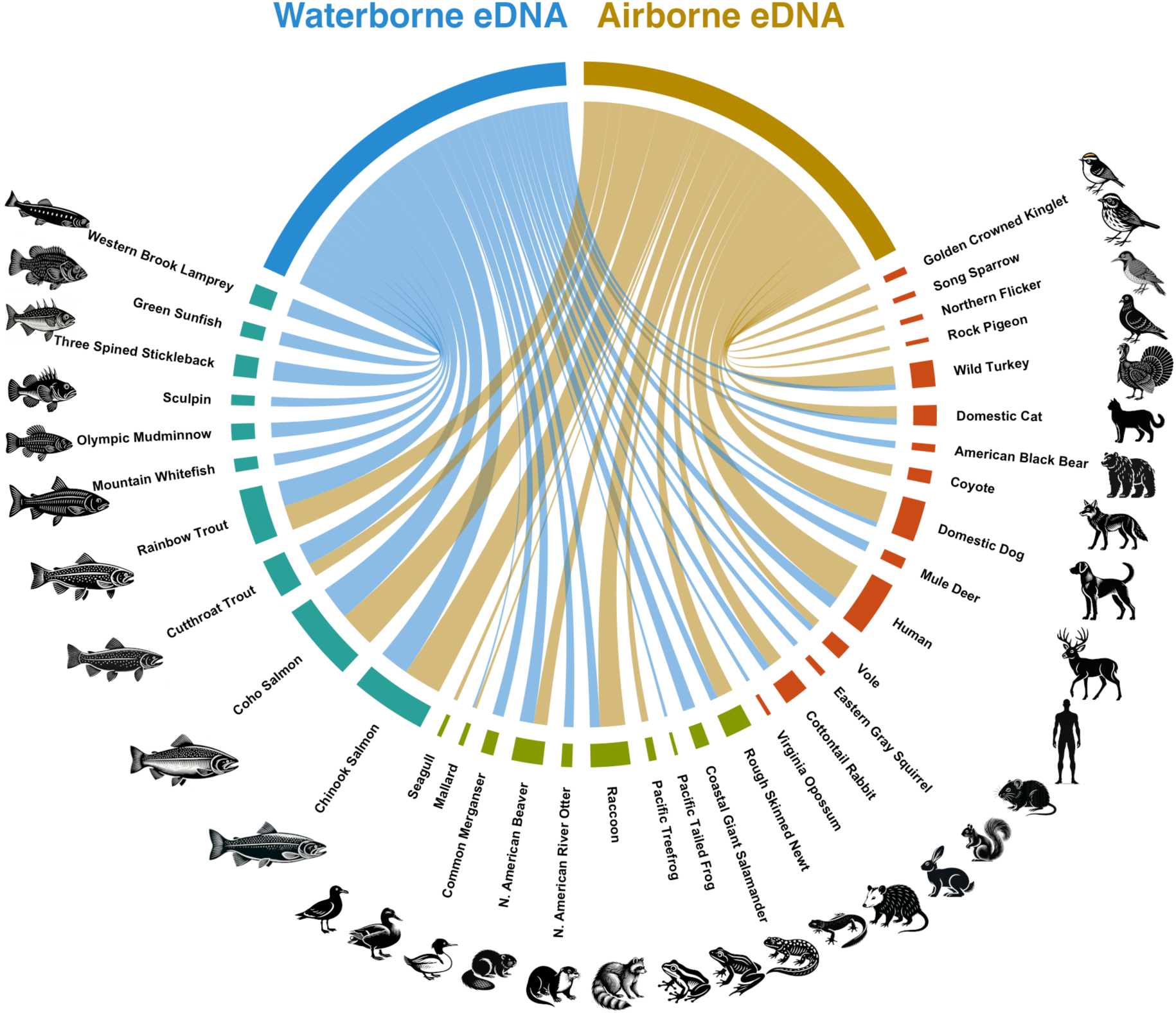
Chord diagram showing overall cross-medium eDNA signal between water and air. Chord ribbons connect the two sampling media—blue arc = water eDNA; gold arc = air eDNA—linked to each detected taxon, with ribbon thickness proportional to the log₁₀-transformed total read count recovered. Outer arc segments are colored by habitat affinity: teal = aquatic, olive = amphibious, and brown = terrestrial.

**Table 1.** Vertebrate taxa sorted by media they were detected in (excluding four “food” species).

| Detection media | Habitat / Biology | Common Name | Species |
| --- | --- | --- | --- |
| Shared | Amphibious | Mallard | *Anas platyrhynchos* |
| Shared | Terrestrial | Domestic Dog | *Canis lupus* |
| Shared | Amphibious | North American Beaver | *Castor canadensis* |
| Shared | Terrestrial | Domestic Cat | *Felis catus* |
| Shared | Terrestrial | Human | *Homo sapiens* |
| Shared | Terrestrial | Wild Turkey | *Meleagris gallopavo* |
| Shared | Terrestrial | Vole | *Microtus* spp. |
| Shared | Terrestrial | Cottontail Rabbit | *Sylvilagus* spp. |
| Shared | Amphibious | Rough-skinned Newt | *Taricha granulosa* |
| Shared | Aquatic | Cutthroat Trout | *Oncorhynchus clarkii* |
| Shared | Aquatic | Coho Salmon | *Oncorhynchus kisutch* |
| Shared | Aquatic | Rainbow Trout | *Oncorhynchus mykiss* |
| Shared | Aquatic | Chinook Salmon | *Oncorhynchus tshawytscha* |
| Shared | Amphibious | Raccoon | *Procyon lotor* |
| Water-only | Amphibious | Coastal (Pacific) Tailed Frog | *Ascaphus truei* |
| Water-only | Aquatic | Sculpin | *Cottus* spp. |
| Water-only | Amphibious | Coastal Giant Salamander | *Dicamptodon tenebrosus* |
| Water-only | Terrestrial | Virginia Opossum | *Didelphis virginiana* |
| Water-only | Aquatic | Three-spined Stickleback | *Gasterosteus aculeatus* |
| Water-only | Aquatic | Western Brook Lamprey | *Lampetra richardsoni* |
| Water-only | Aquatic | Green Sunfish | *Lepomis cyanellus* |
| Water-only | Amphibious | North American River Otter | *Lontra canadensis* |
| Water-only | Amphibious | Common Merganser | *Mergus merganser* |
| Water-only | Aquatic | Olympic Mudminnow | *Novumbra hubbsi* |
| Water-only | Terrestrial | Mule Deer | *Odocoileus hemionus* |
| Water-only | Terrestrial | Eastern Gray Squirrel | *Sciurus carolinensis* |
| Water-only | Aquatic | Mountain Whitefish | *Prosopium williamsoni* |
| Water-only | Terrestrial | American Black Bear | *Ursus americanus* |
| Air-only | Terrestrial | Coyote | *Canis latrans* |
| Air-only | Terrestrial | Northern Flicker | *Colaptes auratus* |
| Air-only | Terrestrial | Rock Pigeon | *Columba livia* |
| Air-only | Amphibious | Gull | *Larus* spp. |
| Air-only | Terrestrial | Song Sparrow | *Melospiza melodia* |
| Air-only | Amphibious | Pacific Treefrog | *Pseudacris regilla* |
| Air-only | Terrestrial | Golden-crowned Kinglet | *Regulus satrapa* |

### 3.2 Cross-medium detection thresholds

For waterborne to airborne eDNA transfers, the model yielded an intercept β₀ = –3.79 and slope β₁ = 1.35 (SE = 0.235, p = 1.08 × 10⁻⁸), corresponding to a 50% detection probability at log₁₀(water_reads + 1) ≈ 2.82 (≈ 661 raw reads). As for airborne to waterborne eDNA transfers, we obtained β₀ = –0.98 and β₁ = 0.844 (SE = 0.339, p = 0.013), with the 50 % threshold at log₁₀(air_reads + 1) ≈ 1.16 (≈ 13.5 raw reads) (Fig. 2).

**Figure 2.**
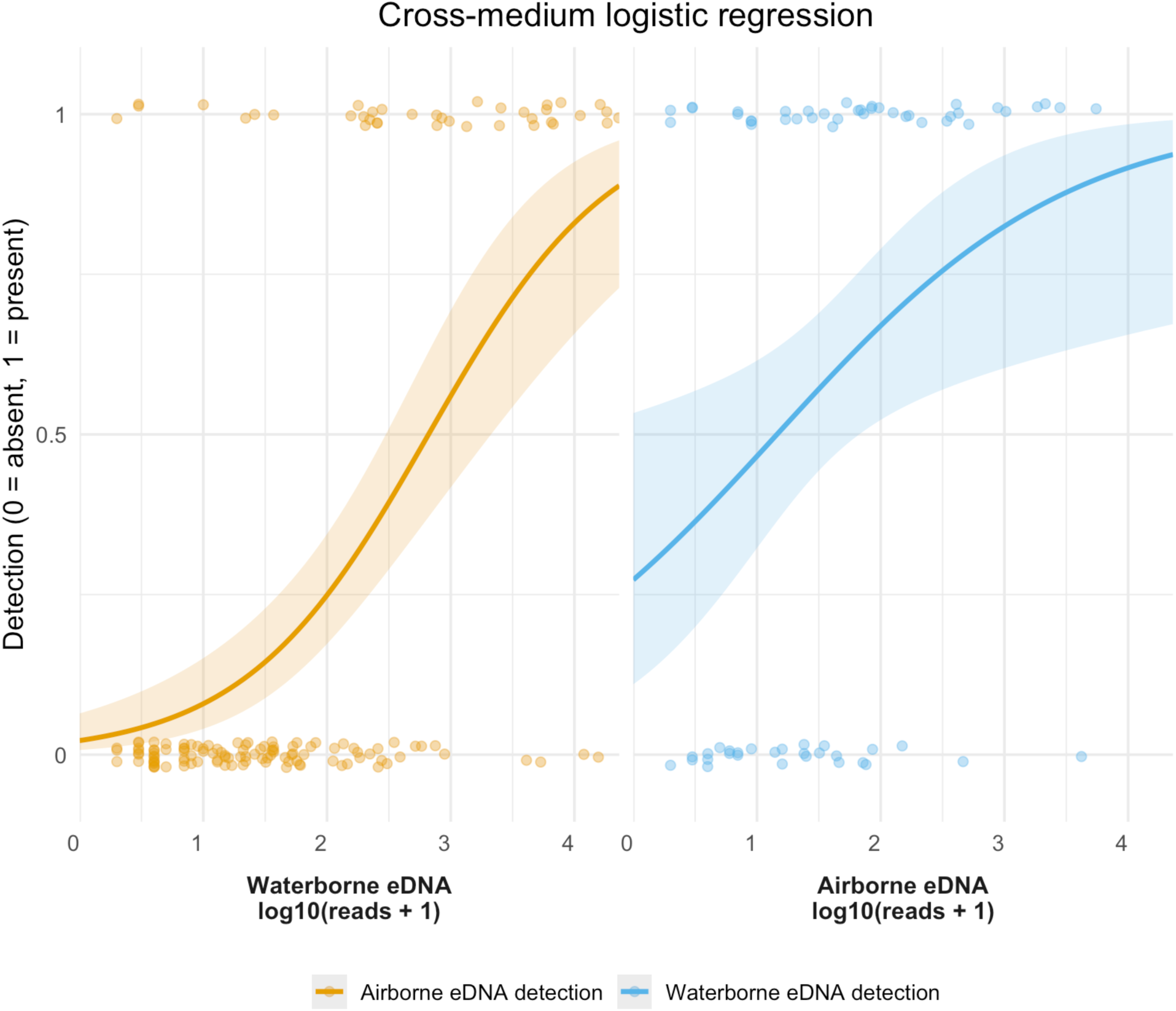
Cross-medium logistic regression of eDNA detection probability. In the left panel, for each species detected in water, golden points indicate whether that species was also detected in air (1 = present, 0 = absent) at each log₁₀-transformed waterborne read count. The golden curve is the fitted binomial GLM, illustrating how higher water eDNA abundance predicts greater probability of exogenous transfer and detection in air. In the right panel, blue points and curve show, for species detected in air, the probability of detecting the same species in water as a function of its airborne eDNA counts. Points are jittered vertically for clarity. Shaded ribbons around each curve represent the 95 % confidence intervals of the logistic regression fit.

The logistic regression shows that once eDNA abundance exceeds medium-specific thresholds, cross-medium detection becomes more predictable, consistent with mechanistic aerosolization and deposition rather than sporadic artifacts. In the water to air regression, fully aquatic taxa (Coho and Chinook Salmon), exhibit a steep slope: once waterborne reads exceed the threshold, the probability of seeing any air detection rises sharply above 50 %. Amphibious species have a much shallower slope and do not reach a 50 % chance of airborne detection at any water-read level. Terrestrial taxa remain at zero transfer probability in the water to air model, suggesting that strictly land-based species detected in the air arise from direct shedding rather than aquatic aerosolization. Conversely, the air to water model shows that aquatic taxa are always detected in water, amphibious taxa rise gradually, and terrestrial taxa require high airborne counts. Together, these patterns confirm that cross-medium detection is fundamentally abundance-driven. A few low-read outliers (e.g. terrestrial DNA in water) reveal genuine “reverse” transfers and emphasize the need to interpret exogenous detections in the context of local ecology and hydrology.

### 3.3 Temporal synchrony across media

Stacked-bar time-series at Issaquah Hatchery (Fig. 3) and Confluence Park (Suppl. Fig. 2) illustrate that the dominant taxa eDNA signals in air and water often peak in the same weeks, even if their exact proportions do not match perfectly. For example, when all taxa are considered (Fig. 3A), salmonids (Chinook and Coho) together comprise > 70% of reads in both media especially in weeks 8 to 10 during the peak spawning run. Pulses of terrestrial detections (e.g., North American Beaver, Mule deer, Wild turkey) also appear in both media at roughly the same times. Focusing on aquatic and amphibious species (Fig. 3B) still shows that salmonid signals rise and fall concurrently in air and water, and even terrestrial and amphibious taxa (Fig. 3C) tend to exhibit their major peaks in both media around the same dates. A few minor deviations, such as a slightly stronger Pacific Treefrog signal in air during Week 11, reflect that low-abundance eDNA signals can decouple briefly since rare taxa detection are inherently unpredictable. Confluence Park (Suppl. Fig. 2) shows a very similar pattern: overall community shifts occur in both air and water at the same general times, but individual proportions sometimes differ.

**Figure 3.**
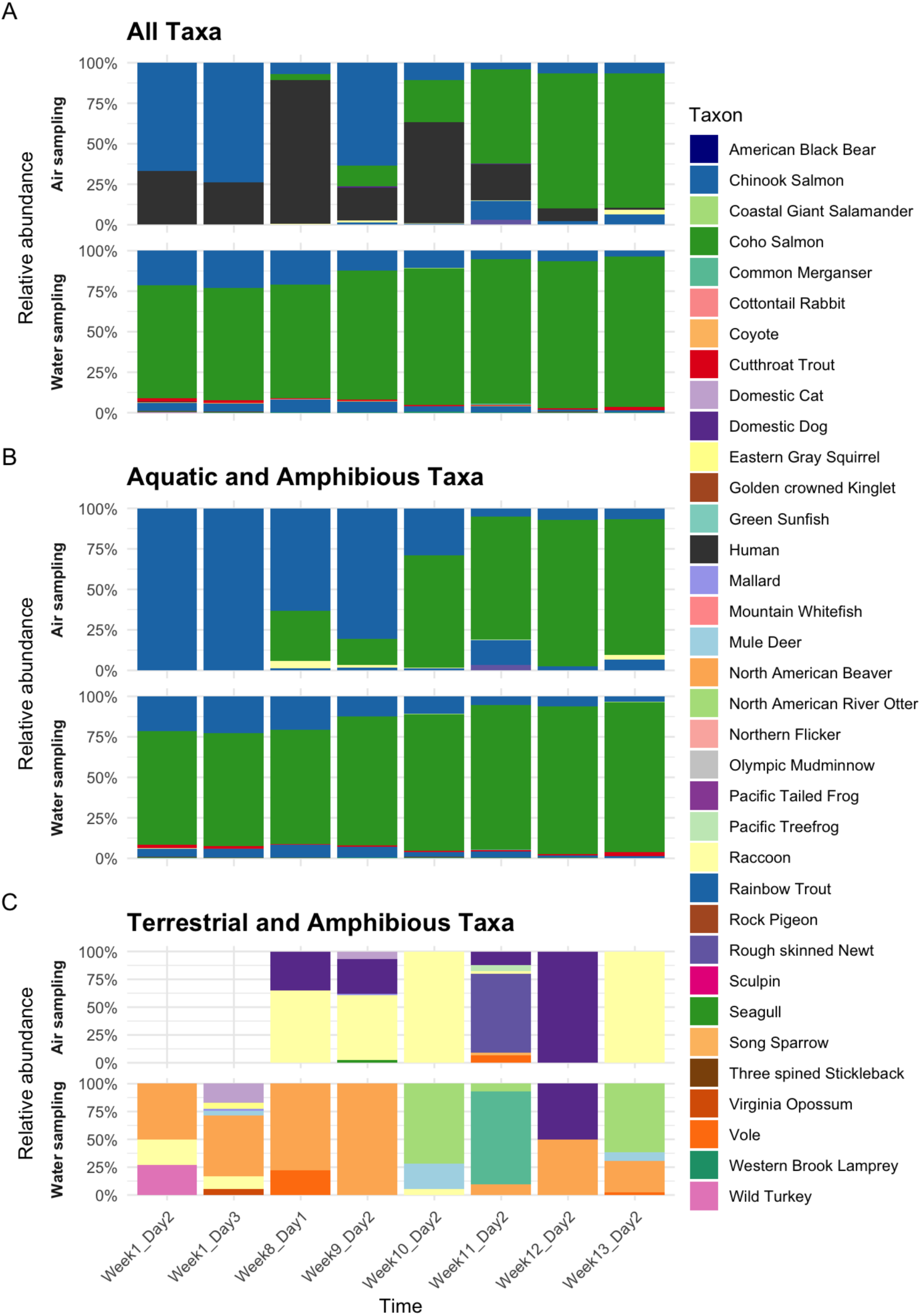
Relative abundance of eDNA-detected taxa at Issaquah Hatchery over time, comparing air versus water sampling. (A) All detected taxa. (B) Only aquatic and amphibious taxa. (C) Only terrestrial and amphibious taxa, omitting humans. In each panel, the top row (Air sampling) and bottom row (Water sampling) show stacked barplots of the proportional (100%) taxon composition over eight weeks. Taxa are keyed by color in the legend to the right.

### 3.4 Focal-species eDNA-index reveals synchronized dynamics

We selected six focal taxa based on their high total read counts and consistent detection in both air and water. These were the Coho Salmon, Chinook Salmon, Rainbow Trout, North American Beaver, Raccoon, and Wild Turkey, thereby representing aquatic, amphibious, and terrestrial guilds. For each species, we computed an eDNA-index (normalized relative abundance scaled 0–1; Kelly et al. 2019, Guri et al., 2024) and plotted its time-series across all sampling dates (Fig. 4).

**Figure 4.**
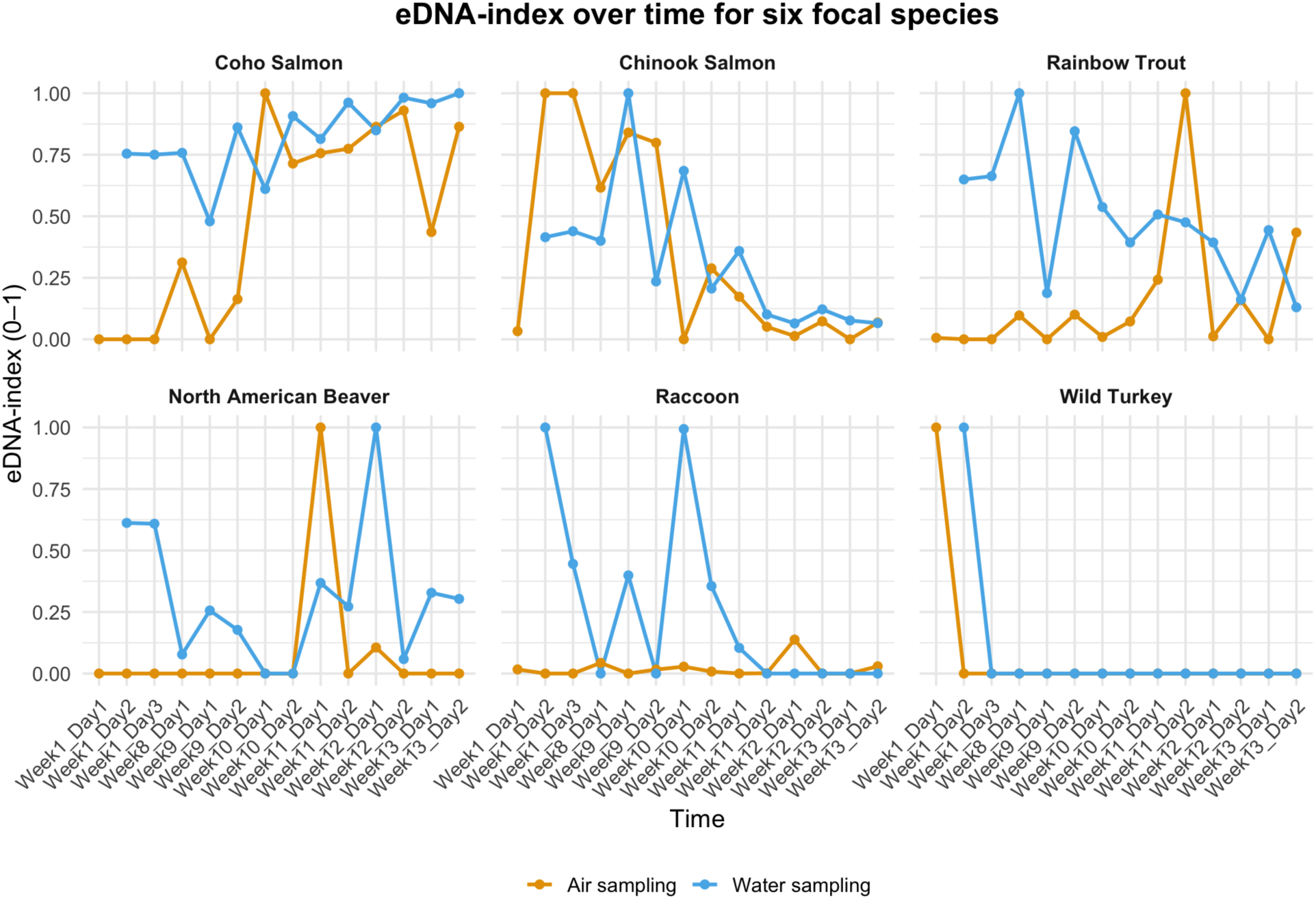
Temporal dynamics of eDNA-index for six focal species. Each panel shows the normalized eDNA index (0–1), across the Time (Week_Day). Gold lines and points represent airborne eDNA sampling; blue lines and points represent waterborne eDNA sampling, observing if both lines match/track one another. Panels (A)–(F) correspond to Coho Salmon, Chinook Salmon, Rainbow Trout, North American Beaver, Raccoon, and Wild Turkey, respectively, illustrating how relative abundance in each medium varies over time. A point at y = 0 indicates a sample was collected but yielded no reads for that species on that date; absence of a point indicates no sample was taken.

Since water samples were collected at the start of each trip immediately before deploying the 24 h passive air traps, any apparent one-day offset between waterborne and airborne peaks reflects our sampling schedule rather than a true biological lag in eDNA transfer. Coho Salmon exhibited a pronounced waterborne peak at Week 8 Day 1 (index ≈ 0.95) with the matching airborne peak recorded in the subsequent 24 h sampling window (Week 8 Day 2). Chinook Salmon reached virtually identical maxima in both media on Week 8 Day 1 (index ≈ 1.0). Rainbow Trout peaked in water at Week 8 Day 2 (index = 1.0) and then produced another airborne pulse at Week 11 Day 1. North American Beaver spiked in water at Week 11 Day 2 (index = 1.0) with a coincident but lower-amplitude air signal on the same date. The amphibious Raccoon (which habitually washes its food at the creekside) displayed an isolated waterborne peak from Weeks 8 to 11 (index ≈ 0.8) that was followed by a brief airborne detection in Week 12. Wild Turkey was detected only at the very first sampling points in Week 1 in both media and was absent thereafter.

Together, these six species’ time-series demonstrate that for the most abundant taxa within our 24 h sampling window, eDNA dynamics in air and water are tightly coupled with minimal temporal offset. This is consistent with synchronous shedding, aerosolization, and deposition, whereas the one-off patterns of lower-biomass guild members (Raccoon, Wild Turkey) underscore the stochasticity of rarer signals. Indeed, when we quantified each species’ air–water concordance (R² from index_air ∼ index_water) and plotted it against the log₁₀ of its total reads (summed across all air + water samples), we found a significant positive trend (β ≈ 0.10 ± 0.03, p = 0.02, adj-R² = 0.48), confirming that more abundant species exhibit stronger synchrony (Suppl. Fig. 3).

### 3.5 Stochastic detection of rare taxa demands sampling effort

Detection frequency, calculated from the proportion of samples in which each taxon appears, varies strongly with total read count (scaled by √ –square root point size; Fig. 5). The most abundant species (e.g. Chinook Salmon, Coho Salmon) were detected in >90% of both airborne and waterborne samples, whereas low-abundance taxa (e.g. Mallard, Pacific Treefrog) fell below 10% in either medium. The tight, positive relationship between √(total reads) and detection probability implies that reliably capturing rare signals would likely require several-fold more replicates than for dominant taxa.

**Figure 5.**
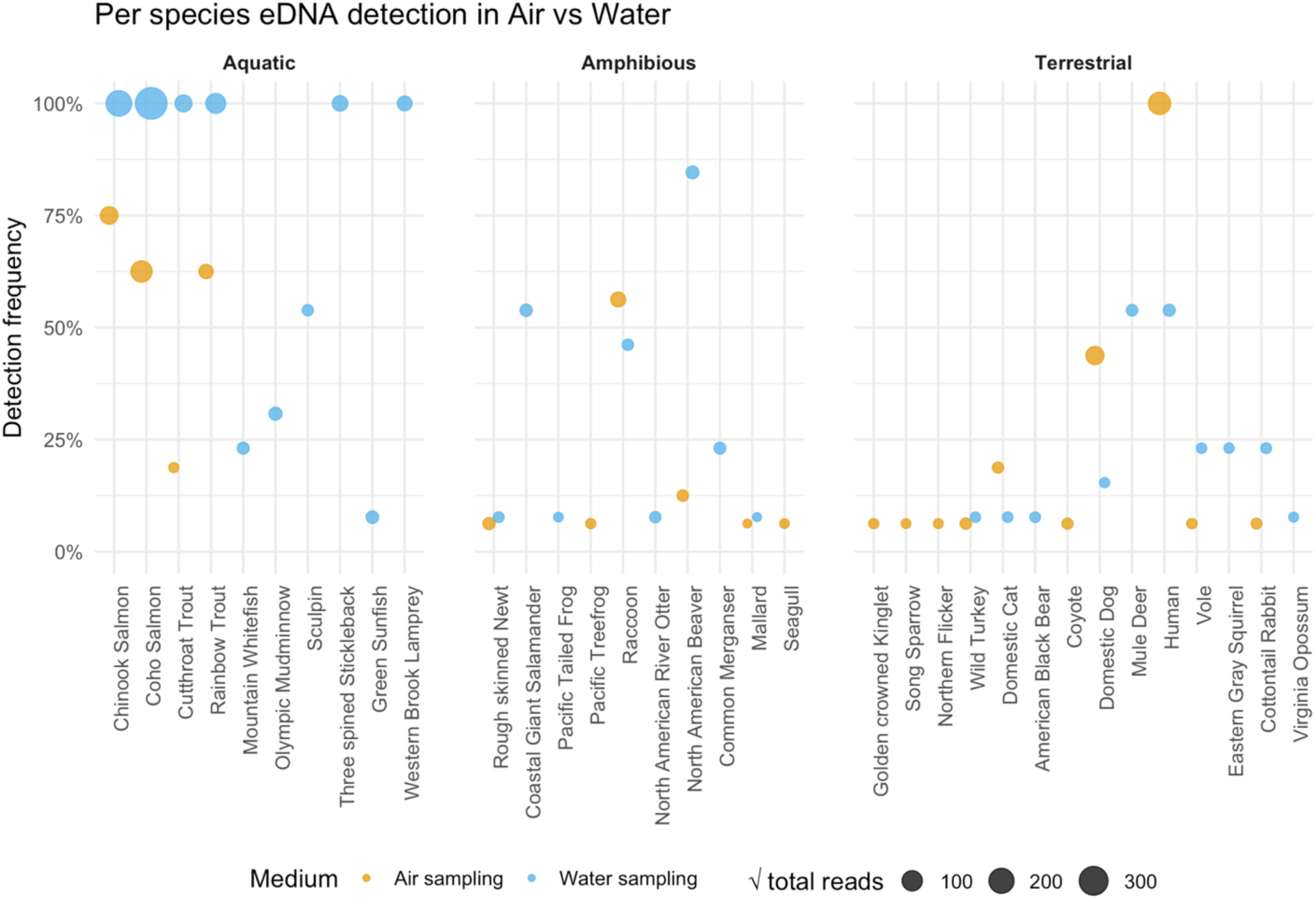
Detection frequency of eDNA for individual species in airborne vs. waterborne samples, grouped by habitat. Each point shows the percentage of samples in which a given species was detected (y-axis) in air (gold) or water (blue), with point size proportional to the square-root of the total read count. Species are ordered and faceted by habitat category (Aquatic, Amphibious, Terrestrial) to illustrate how detection success varies by medium and ecological guild.

### 3.6 Medium drives community structure

Non-metric multidimensional scaling of both presence–absence (Jaccard; stress = 0.126) and abundance (Bray–Curtis; stress = 0.173) matrices (Fig. 6A–F) revealed that samples cluster by sampling medium (air vs. water). In the Jaccard ordination of all taxa (Fig. 6A), air and water points form two distinct clouds, and the Bray–Curtis ordination (Fig. 6D) shows the same pattern. Subsetting to aquatic and amphibious taxa (B, E) or terrestrial and amphibious taxa (C, F) yields similar medium-driven separation. Permutational ANOVA (adonis2, 999 permutations) on Jaccard distances showed that medium (air vs. water) explained the largest share of variance (R² ≈0.32, p=0.001), with location and week contributing small but significant effects (R² ≈0.05– 0.06, p<0.05), and day having no effect. On Bray–Curtis distances, medium again dominated (R² ≈0.44, p=0.001), week was significant (R² ≈0.07, p=0.009), and location/day were non-significant (p>0.05). Although air– and water-derived communities cluster distinctly, a sizable proportion of shared taxa and predictable cross-medium transfers reveals that they nevertheless function as a unified eDNA continuum.

**Figure 6.**
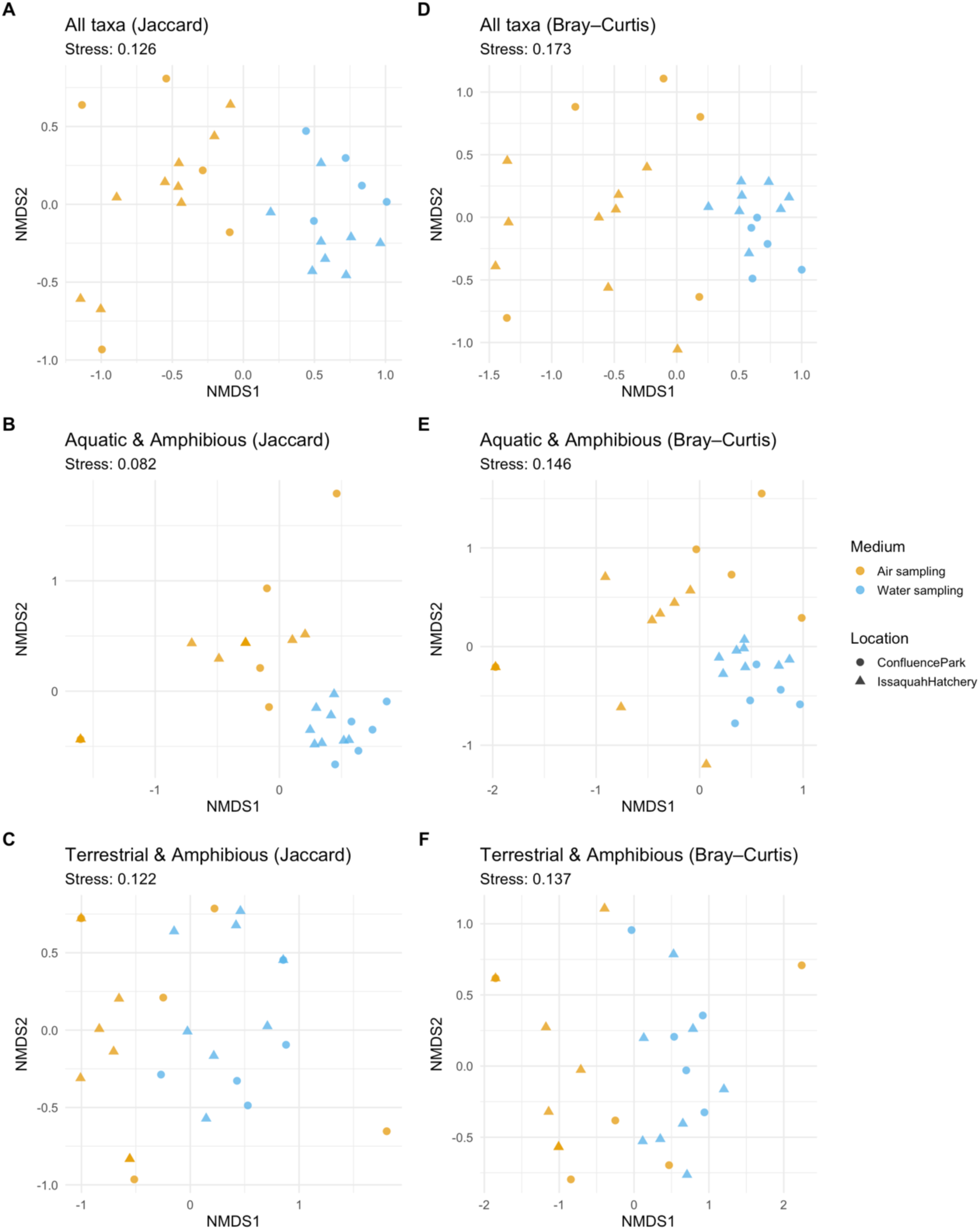
nMDS of eDNA community composition across sites and sampling media using Jaccard dissimilarity (left) and Bray-Curtis similarity (right). Points represent individual samples with color indicating sampling medium (gold = air, blue = water) and shape indicating site (circle = Confluence Park, triangle = Issaquah Hatchery). Stress values for each analysis are shown beneath the panel titles. (A) All taxa: nMDS based on presence/absence of all detected taxa. (B) Aquatic + Amphibious taxa: nMDS restricted to taxa classified as aquatic or amphibious. (C) Terrestrial + Amphibious taxa: nMDS restricted to taxa classified as terrestrial or amphibious.

## 4. Discussion

### 4.1 An Authentic Air-Water Biodiversity Continuum

Our side-by-side survey of river-grab filters and passive open-water trays provides the first field-scale confirmation that water and air form a single, intertwined reservoir of eDNA. While water and air samples offer partially complementary signals, aquatic and airborne eDNA co-occur for at least 40% of detected taxa, ranging from salmon (*Oncorhynchus* spp.) to raccoons (*Procyon lotor*), and at rates exceeding random expectation. This “genetic Ferris Wheel” mirrors natural processes: stream turbulence and bubble-burst events likely help to aerosolize waterborne DNA; rainfall and overland flow return terrestrial signals to the river; and gravitational settling deposits airborne fragments onto surfaces (Allen et al., 2022; Evangeliou et al. 2020; Xu and Nukazawa, 2025; Zinger et al., 2025). By revealing these linked pathways, our work breaks down the traditional water-versus-air compartments of eDNA research and calls for a unified, continuum-based perspective on genetic monitoring (Deiner et al., 2016; Lynggaard et al., 2022; Ip et al., 2025a). Building on the salmon-focused qPCR work of Ip et al. (2025a), which first showed that spawning salmon release detectable DNA into the air, we here broaden that insight to a fuller vertebrate community. Progressing from targeting a single species, we metabarcoded 35 taxa, and rather than sampling air alone, we paired open-water trays with river grab filters, enabling us to derive generalizable abundance-driven thresholds for cross-medium transfer.

Importantly, this continuum is not random background noise but follows clear, abundance-driven rules. Because abundant DNA molecules in one compartment will always feed the next, high eDNA concentration in water inevitably produces some airborne fragments, and vice versa. Our models identify 50 % detection probability thresholds of approximately 660 raw water reads to predict airborne detection, and about 14 raw air reads to predict waterborne detection. As each water sample (1 L) and each air sample (750 cm² × 24 h) represent very different sampling substrates—and we did not directly measure airborne deposition to convert into an equivalent air volume—these thresholds must be interpreted qualitatively rather than as absolute concentration comparisons. Future work should quantify deposition rates under comparable field conditions to normalize air-versus-water sampling effort. In the water-to-air model, fully aquatic taxa, especially those engaged in high-energy behaviors like spawning that generate bubble-burst aerosols, exhibit the steepest curves, reflecting their efficient transfer of DNA from water into air. Amphibious species, by virtue of using both aquatic and terrestrial habitats, will shed DNA into each medium directly. As a result, their air-detection curves represent a combination of water-to-air transfer and direct terrestrial shedding, yielding intermediate slopes relative to fully aquatic or strictly terrestrial taxa. Truly terrestrial species (for example, deer and wild turkey) fall outside this water-to-air framework: their DNA never passes through the aquatic pool but is released directly into the atmosphere via hair, skin cells, or feces, producing essentially flat dose–response curves when plotted against water read counts. To capture their dynamics, we instead employ an air-to-water model in which airborne abundance predicts the occasional appearance of land-derived DNA in water through settlement, deposition, runoff, rain-splash, or overland flow.

By fitting two targeted GLMs, one for water-to-air transfers among aquatic and amphibious taxa, and one for air-to-water transfers among terrestrial taxa, we account for each group’s unique shedding and transport pathways and avoid forcing a single model to fit all eco-hydrological-to-atmospherical contexts. By linking our study-specific read-count benchmarks to dispersal mechanics, we illustrate how practitioners could, in principle, plan sampling regimes to hit a desired detection probability—while noting that these thresholds (≈660 water reads, ≈14 air reads) will vary with community composition, sequencing depth, and lab protocols and thus should be calibrated for each new system. By spanning taxa from salmon to amphibians, birds and mammals, our results fulfill the conceptual ‘Air-Water eDNA Continuum’ framework, demonstrating its applicability across diverse vertebrate groups.

### 4.2 Temporal Synchrony and Stochastic Boundaries

If water and air truly share DNA, their temporal patterns should echo one another, and indeed, our time-series data aligned well. Peaks in Coho and Chinook salmon eDNA in water were mirrored in passive air traps over the same 24 h sampling window (i.e., air traps were deployed immediately after each water grab), while North American Beaver and Raccoon produced concordant but more gradual rises and falls. This synchrony confirms that shedding, aerosolization, and deposition operate on comparable timescales across media, enabling airborne eDNA to serve as a faithful, near–real-time sentinel of aquatic population pulses. Occasional anomalies such as an isolated Raccoon eDNA pulse in air with no matching water signal, likely reflect discrete terrestrial shedding events or weather-driven dispersal, but these do not undermine the overall alignment. By contrast, Wild Turkey eDNA was recovered in both air and water only during the first sampling event on consecutive days, after which it never reappeared in either medium, highlighting how low-abundance terrestrial signals may flicker at the very edge of the continuum. Importantly, the degree of synchrony itself scales with abundance (see end of Section 3.4 and Supplementary Fig. 3), confirming that more abundant species exhibit tighter air–water coupling while rarer taxa remain stochastic.

Despite the overall synchrony, this continuum frays at low abundances. While metabarcoding read counts are not perfect measures of absolute biomass, our experimental design’s intentional absence of post-PCR normalization means that raw reads here track relative abundance closely enough to support these thresholds. Our detection-frequency analysis shows that taxa yielding on the order of 10 reads or fewer appear erratically, with detection rates often below 10%. When targeting rare taxa, the data clearly show that exogenous cross-medium detections become effectively random. In these cases, it is far more efficient to concentrate sampling effort in the medium where the species is expected; intensively replicating water grabs for aquatic taxa or air traps for terrestrial taxa; rather than chasing sporadic spillover signals. For rare or low-biomass taxa, reliable monitoring requires scaled-up replication, typically three to five times more traps or water grabs to overcome stochastic “flicker” and achieve desired confidence levels. Recognizing these stochastic boundaries is essential for conservation applications targeting elusive or endangered species, where effort must be matched to the expected eDNA yield to avoid false negatives.

Beyond these practical thresholds, our data hint at richer dynamics yet to be explored. Quantifying time-lagged cross-correlations between water and air eDNA for different taxa could reveal species-specific transport kinetics—how quickly each organism’s DNA moves through the continuum under varying flow regimes, wind conditions, or diel activity cycles. Such insights would allow adaptive sampling schedules, timed to coincide with peak shedding or optimal transport conditions, further sharpening the power of bidirectional monitoring.

Finally, taxa yielding on the order of 10 reads or fewer exhibited detection frequencies below 10% across samples, providing a practical rule of thumb for our dataset but one that may vary with sequencing depth, primer choice, and local eDNA concentrations. Strong, abundant eDNA signals produce near-immediate cross-medium echoes, whereas rare signals demand targeted, medium-specific sampling coupled with intensified replication. By mapping these stochastic boundaries, we equip practitioners with the knowledge to deploy continuum-based surveys both efficiently and effectively.

### 4.3 Applied Frontiers and Near-Term Outlook

Beyond vertebrate eDNA, similar aerosolization and deposition processes have been documented for microplastics (Allen et al., 2022; Evangelou et al., 2024; Evangeliou et al. 2020), soil eDNA in rain-wash events (Xu & Nukazawa, 2025), and airborne microbial DNA recovered over Antarctic deserts (Bottos et al., 2014). The empirical framework we establish opens multiple avenues for immediate, impactful applications. First, the passive open-water trays require no specialized equipment, just a plastic tray with deionized or mineral water (Klepke et al., 2022), making them ideal for remote, resource-limited settings where traditional sampling is impractical. Managers can deploy “passive deposition samplers” in riparian corridors to generate holistic snapshots of vertebrate communities, democratizing biodiversity monitoring by engaging citizen scientists including hikers, tribal monitors, and school groups to participate in cross-medium campaigns.

Second, the continuum paradigm underpins powerful early-warning systems (Goldberg et al., 2015). Paired air–water eDNA deployments could flag the first arrivals of non-native taxa before they are caught by visual surveys or camera traps. (Jerde et al., 2013). Likewise, restoration projects reintroducing fish or amphibians into degraded waterways could leverage airborne eDNA as a rapid, noninvasive sentinel of colonization success (Evans et al., 2017), detecting massive breeding-driven eDNA pulses in both water and air and providing real-time feedback on project outcomes (Tillotson et al., 2018; Ip et al., 2023).

Third, the bidirectional framework extends naturally to One Health surveillance. Pathogenic microbes and viruses should follow the same cross-medium dynamics: waterborne pathogens aerosolized by bubble bursts or rainfall splashes may become detectable in air, while airborne pathogens settling onto water surfaces could signal emerging contamination events (Blanchard and Syzdek, 1970; Gholipour et al., 2021). Paired sampling thus offers a unified sensor network for zoonotic and waterborne threats at wildlife–human interfaces, enhancing early detection and response capabilities (Lednicky et al., 2020).

Finally, our abundance-driven thresholds and temporal synchrony lay the groundwork for predictive eDNA modeling. By integrating read-count benchmarks with particle-size distribution data, hydrological measurements, aerosol physics, and eDNA decay kinetics (Brandão-Dias, et al., 2025; Fröhlich-Nowoisky et al., 2016; Jane et al., 2015), one can forecast cross-medium hotspots much like air-quality models predict pollutant plumes. Coupled with real-time meteorological and flow sensors (Sheik et al., 2024), such predictive tools could power dynamic eDNA dashboards, showing automated alerts guiding adaptive sampling and triggering rapid management interventions in the face of biodiversity pulses or contamination events.

Embedding these samplers into existing monitoring networks, managers can move from retrospective inventories to real-time biodiversity intelligence. By anchoring the Bidirectional eDNA Continuum Paradigm in field data, we catalyzed a conceptual shift in how we monitor, model, and manage life on Earth. Water and air are no longer separate chapters in the genetic manuscript of biodiversity; they are two pages of the same story, bound together by physical processes and biological behaviors. As environmental genomics scales to address global conservation, biosecurity, and public health challenges, embracing this unified reservoir and the genetic Ferris-wheel will be essential for capturing the full breadth, tempo, and connectivity of living systems.

## 5. Acknowledgements

We thank Natasha Kacoroski, Larry Franks, and the dedicated volunteers at Friends of the Issaquah Salmon Hatchery for their generous support in the field. We are also grateful to Travis A. Burnett and Darin Combs at the Washington Department of Fish and Wildlife for facilitating access and allowing field experimental work at the Issaquah Hatchery. We acknowledge funding support from the National Philanthropic Trust and the Packard Foundation [Grant No. 2021-72609] and thank the Center for Environmental Genomics at the University of Washington for providing access to high-performance computing resources.

## Author Contributions

Y.C.A.I. conceived the study. Y.C.A.I. and E.A.A. designed the field and laboratory protocols. Y.C.A.I. and P.B.D.F.P., jointly designed the downstream bioinformatic analyses. P.B.D.F.P. conducted the bioinformatic analyses, while Y.C.A.I, G.G. and P.B.D.F.P. conducted the statistical analyses with inputs from R.P.K. The fieldwork was performed by Y.C.A.I. and E.A.A., while Y.C.A.I. wrote the manuscript. R.P.K. supervised the project, contributed to conceptual guidance, and provided critical revisions. All authors contributed to the study design and approved the final manuscript.

Data and Materials Availability:

The authors declare that they have no competing interests. All data needed to evaluate the conclusions in this paper are available in the main text and/or the Supplementary Materials. Additional data, code, and materials will be made available upon reasonable request. No materials were subject to material transfer agreements (MTAs).

## 7. Supplementary Figures and Table

**Supplementary Figure 1.**
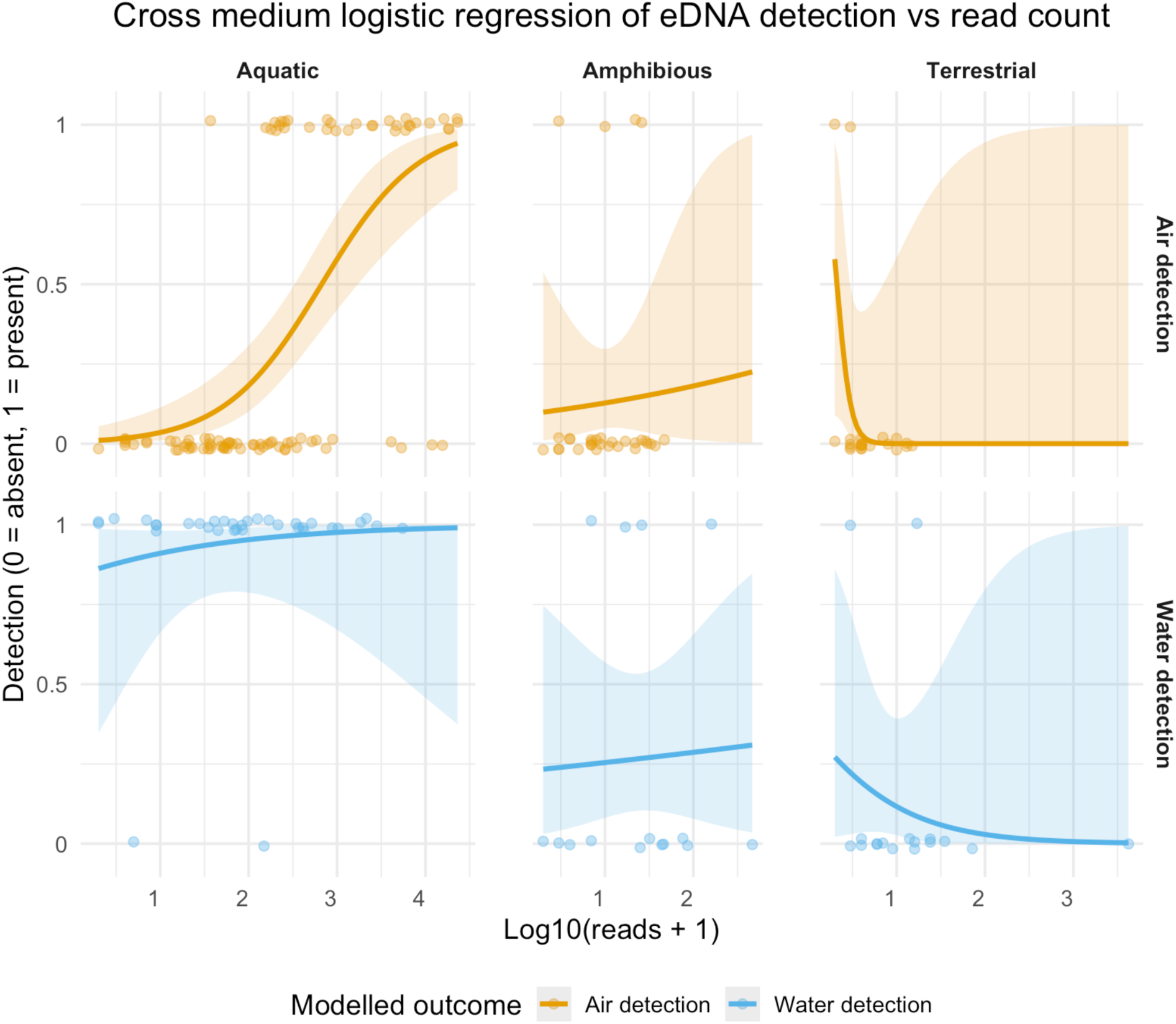
Cross-medium logistic regression of eDNA detection probability by ecological guild. Each column shows one habitat guild; Aquatic, Amphibious, and Terrestrial, and each row shows detection in the alternate medium: Air detection (top row) and Water detection (bottom row). On the x-axis is the log₁₀-transformed read count (reads + 1) in the source medium; on the y-axis is the binary detection outcome in the target medium (1 = present, 0 = absent). Gold points and curves correspond to the probability of detecting a taxon in air as a function of its waterborne eDNA abundance; blue points and curves correspond to the probability of detecting a taxon in water as a function of its airborne eDNA abundance. Points are jittered vertically for clarity, and the smooth lines are fitted binomial GLMs. Shaded ribbons around each curve represent the 95 % confidence intervals of the logistic-regression fit.

**Supplementary Figure 2.**
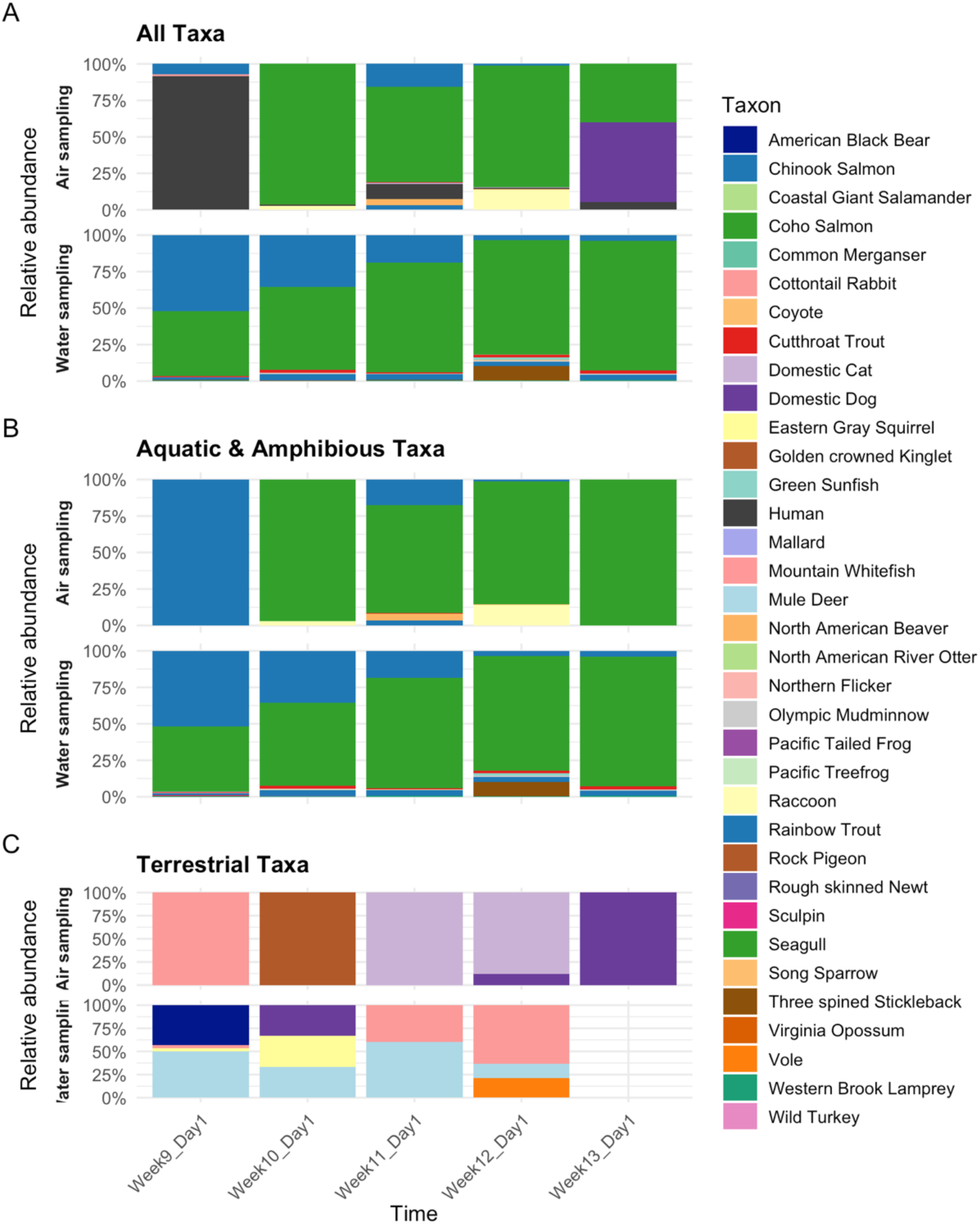
Relative abundance of eDNA-detected taxa at Confluence Park over time, comparing air versus water sampling. (A) All detected taxa. (B) Only aquatic and amphibious taxa. (C) Only terrestrial taxa. In each panel, the top row shows air-sampling filter results and the bottom row shows water-sampling filter results, with stacked bars representing the proportional (100%) taxon composition over five weeks. Taxa are keyed by color in the legend to the right.

**Supplementary Figure 3.**
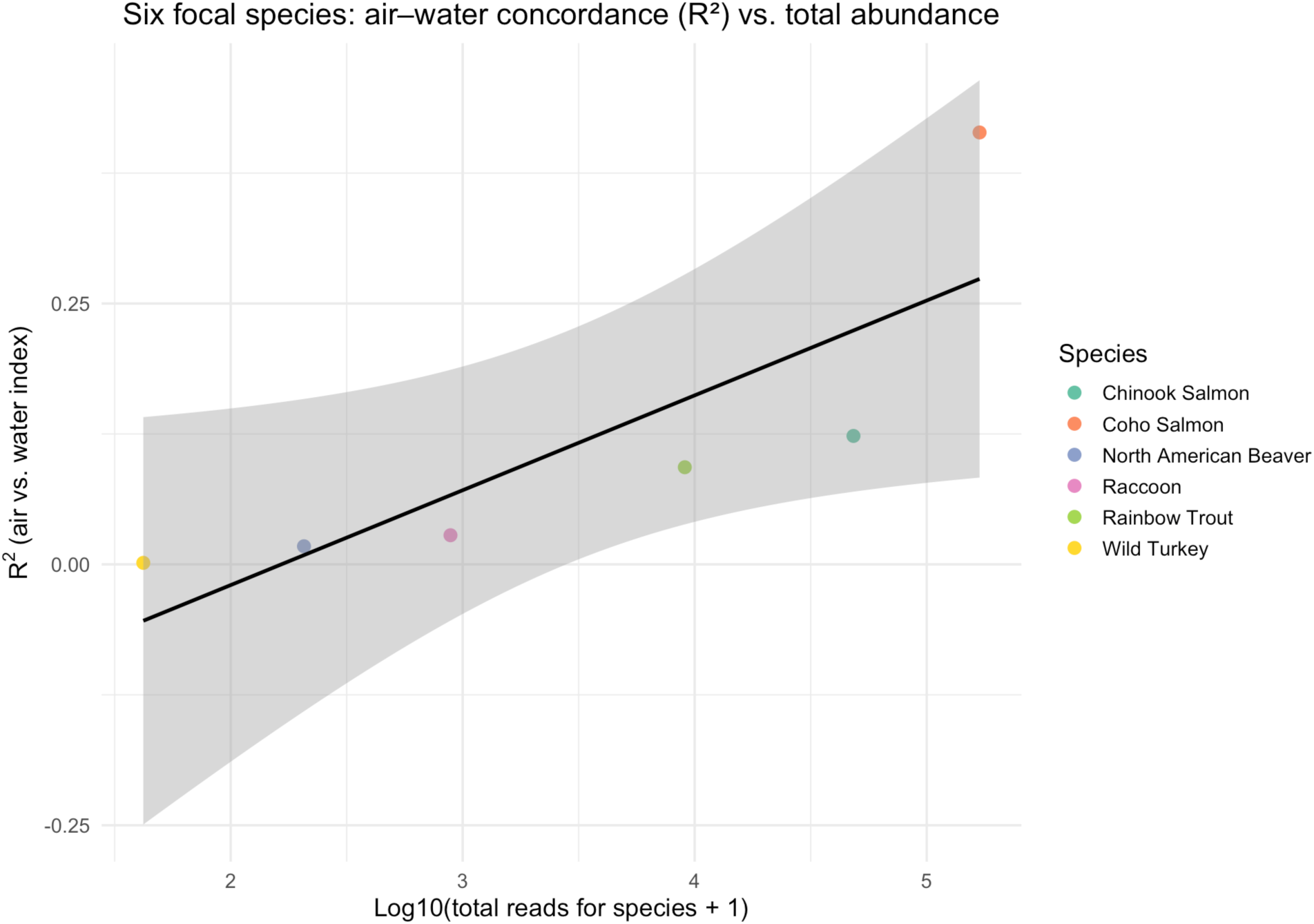
Species-level air–water concordance (R²) as a function of overall eDNA read abundance. Each point represents one of the six focal taxa (colored by species), plotting its R² from a linear regression of air-index versus water-index against log₁₀(total reads + 1) across all samples. The solid black line is the best-fit linear model (β ≈ 0.10 ± 0.03 SE, p = 0.02, adj-R² = 0.48), with the shaded ribbon showing the 95% confidence interval.

**Supplementary Table S1.** Sequencing Yield Summary for Paired Airborne and Waterborne eDNA Samples.

| SampleCode | Location | PreFilterReads | PostFilterReads |
| --- | --- | --- | --- |
| Air018 | IssaquahHatchery | 22774 | 12764 |
| Air024 | IssaquahHatchery | 215 | 97 |
| SAL042 | IssaquahHatchery | 23532 | 7672 |
| Air025 | IssaquahHatchery | 77 | 48 |
| SAL049 | IssaquahHatchery | 58793 | 17129 |
| Air036 | IssaquahHatchery | 8481 | 4865 |
| SAL093 | IssaquahHatchery | 117360 | 32543 |
| SAL149 | ConfluencePark | 39648 | 8787 |
| Air042 | ConfluencePark | 1749 | 1208 |
| SAL101 | ConfluencePark | 141255 | 35081 |
| Air048 | IssaquahHatchery | 10543 | 3361 |
| SAL105 | IssaquahHatchery | 67800 | 20293 |
| Air054 | ConfluencePark | 26740 | 5710 |
| SAL113 | ConfluencePark | 51333 | 11526 |
| Air060 | IssaquahHatchery | 4464 | 1975 |
| SAL117 | IssaquahHatchery | 28452 | 7184 |
| Air066 | ConfluencePark | 1530 | 620 |
| SAL125 | ConfluencePark | 129170 | 24410 |
| Air072 | IssaquahHatchery | 9770 | 3171 |
| SAL129 | IssaquahHatchery | 22410 | 5289 |
| Air078 | ConfluencePark | 16636 | 3387 |
| SAL137 | ConfluencePark | 30601 | 7546 |
| Air084 | IssaquahHatchery | 4571 | 1236 |
| SAL141 | IssaquahHatchery | 53746 | 12248 |
| Air090 | ConfluencePark | 152 | 130 |
| Air096 | IssaquahHatchery | 3179 | 1080 |
| SAL153 | IssaquahHatchery | 120680 | 24892 |

Supplementary Material 1. Codes for post demultiplexing sequence processing

